# MutationAssessor in cBioPortal

**DOI:** 10.1101/2025.08.10.669566

**Authors:** Yang Su, Xiang Li, Boris Reva, Yevgeniy Antipin, Nikolaus Schultz, Ino de Bruijn, Chris Sander

## Abstract

MutationAssessor (MA) helps researchers evaluate the likely functional impact of somatic and germline mutations in cancer. It provides an evolution-based functional impact score (FIS) to classify mutations based on their likely effect on protein function. FIS scores are based on analysis of patterns of conservation in protein families (conserved residues) and subfamilies (specificity residues). In this new version (r4) we have (1) refined the combinatorial entropy analysis of conservation patterns, (2) recalculated full-length protein multiple sequence alignments covering a larger fraction of human proteins and making use of the explosive growth of protein sequence data, (3) compared predicted functional impact with the pathogenic-benign classification of sequence variants in curated knowledge bases, such as ClinVar, (4) observed the inverse relationship between predicted high functional impact and variant frequency in germline genome sequences and (5) explore the evaluation of switch-of-function mutational effects. Functional impact of ∼4 million somatic amino-acid changing mutations across more than 320K human tumor samples are now available in the widely used cBioPortal for Cancer Genomics.

## Introduction

The Cancer Genome Atlas (TCGA) and related projects at major cancer centers have generated large datasets on genomic variants in cancer tissues, including amino acid (AA)-changing somatic mutations that affect the function of particular proteins, such as oncogenes and tumor suppressors (AACR Project GENIE Consortium, 2017; Kandoth et al., 2013). The widely used cBioPortal for Cancer Genomics makes these data available for cancer researchers (de Bruijn et al., 2023). cBioPortal is an open-access platform designed for visualizing, analyzing, and exploring large-scale cancer genomics datasets. It integrates diverse molecular data, including mutations, copy number alterations, gene expression, and clinical data. The portal allows interactive exploration of patient cohorts, helping researchers identify cancer-related genomic alterations, oncogenic pathways, and correlations of variants with clinical outcomes. It also provides custom analysis tools, including survival plots, mutation frequency comparisons, mutation hotspots (Gao et al., 2017; Gauthier et al., 2016; Miller et al., 2015), and co-occurrence analysis. Widely used in precision oncology and translational research, cBioPortal accelerates discoveries by making complex cancer genomics data accessible (e.g., for clinical interpretation of personalized variant profiles) and actionable (for the design of experimental and computational cancer biology research projects). One of the mutation effect tools in cBioPortal is MutationAssessor (Reva et al., 2011).

Various computational tools have been developed to predict the functional impact of amino-acid changing variants, including well-established methods such as SIFT (Ng and Henikoff, 2003), PolyPhen-2 (Adzhubei et al., 2010), MutationTaster (Schwarz et al., 2014), CADD (Rentzsch et al., 2019), REVEL (Ioannidis et al., 2016) and GEMME (Abakarova et al., 2023). These tools span a spectrum of approaches, from conservation-based scoring to supervised training on known mutation effect data and majority-vote ensemble models. MutationAssessor falls into the class of unsupervised, conservation-based tools, uniquely incorporating subfamily-specific conservation to detect functional shifts that global conservation metrics may miss (Gnad et al., 2013). Its strengths lie in interpretability, proteome-wide coverage, and a focus on somatic cancer mutations. While other variant effect prediction tools typically are evaluated in tests against systematic mutational scan assays (Notin et al., 2023), almost all of these assays are conducted *in vitro* under specific conditions and are not assured to reflect functional relevance in physiological or disease contexts. Instead, MutationAssessor is based on evolutionary constraints on sequence variation in real organisms (Rochman et al., 2020), prioritizes scoring for real-world cancer mutations and makes these predictions easily accessible through integration with the cBioPortal for Cancer Genomics.

Specifically, MutationAssessor uses a cross-species alignment-based method to evaluate positional conservation patterns and to predict the effect of amino acid substitutions on protein fitness, based on the notion that protein fitness contributes to selection in evolution. Aligned sequences of homologs from a protein family are clustered into subfamilies by combinatorial entropy optimization (CEO) (Reva et al., 2007). For an amino acid change in a protein, the resulting combinatorial entropy changes are evaluated in the context of all homologs (conservation score) and in the context of the subfamily to which the wildtype protein belongs (specificity score). The conservation and specificity scores reflect how a mutation is consistent with or violates the conservation patterns observed in evolution within a protein family and its subfamilies. A combination of conservation and specificity scores—the functional impact score (FIS)—predicts the effect of the amino acid change on protein function (see Methods for details). Proteins containing a mutation with positive FIS are predicted to be less fit than the normal (‘wild type’) version; and those with negative FIS are predicted to be more fit than the wild type. The most plausible interpretation is that ‘less fit’ reflects reduced or complete loss of function, while ‘more fit’ may imply gain of function.

MutationAssessor was first released in 2011 (MA r1) (Reva et al., 2011) and subsequently updated (unpublished) in 2012 (MA r2) and 2015 (MA r3). Here we report an improved release, MA r4, with updated functional impact scores for nearly all AA-changing (aka missense) single substitution variants in human proteins, for which reasonably rich evolutionary multiple sequence alignments can be obtained. We make available source code in the public domain, multiple sequence alignments for human proteins via [Genome Nexus](https://www.genomenexus.org/) (de Bruijn et al., 2022), and detailed mutational effect scores in the user-friendly and widely used cBioPortal for Cancer Genomics (cbioportal.org). cBioPortal, originally developed in the context of The Cancer Genome Atlas now covers data from more than 320K tumor samples and a suite of analysis tools and is used by more than 40K unique active users as of early 2025. We describe the latest improvements, updates and user access of MA for cancer researchers; and compare in detail the computed functional impact of sequence variants in well-known oncogenes and tumor repressors with occurrence frequencies of somatic variants across thousands of human tumor samples from The Cancer Genome Atlas (TCGA); and, assess the relationship between high FIS scores and reported pathogenic impact from ClinVar, a curated database of known disease-associated variants.

## Results

### Dual conservation analysis

The power of MutationAssessor is based on two types of conservation analysis across multiple species (Fig. 1). For each human protein in UniProtKB, we build a multiple sequence alignment (MSA) from the amino acid sequence of its canonical isoform. We then cluster the sequences in the MSA into subfamilies using the combinatorial entropy optimization (CEO) method (Reva et al., 2007). Subfamilies are clusters of proteins. The CEO method defines subfamily decomposition of an MSA as the one that optimizes specificity patterns of a selected set of highly-ranked specificity residues: overall variation, but conservation within subfamilies. Definition of the set of specificity residues and of the subfamily organisation is the result of a joint optimization problem, not done in separate steps. The effect of a single-amino acid substitution mutation is then evaluated by applying conservation analysis to both the whole protein family and individual subfamilies.

**Figure 1.**
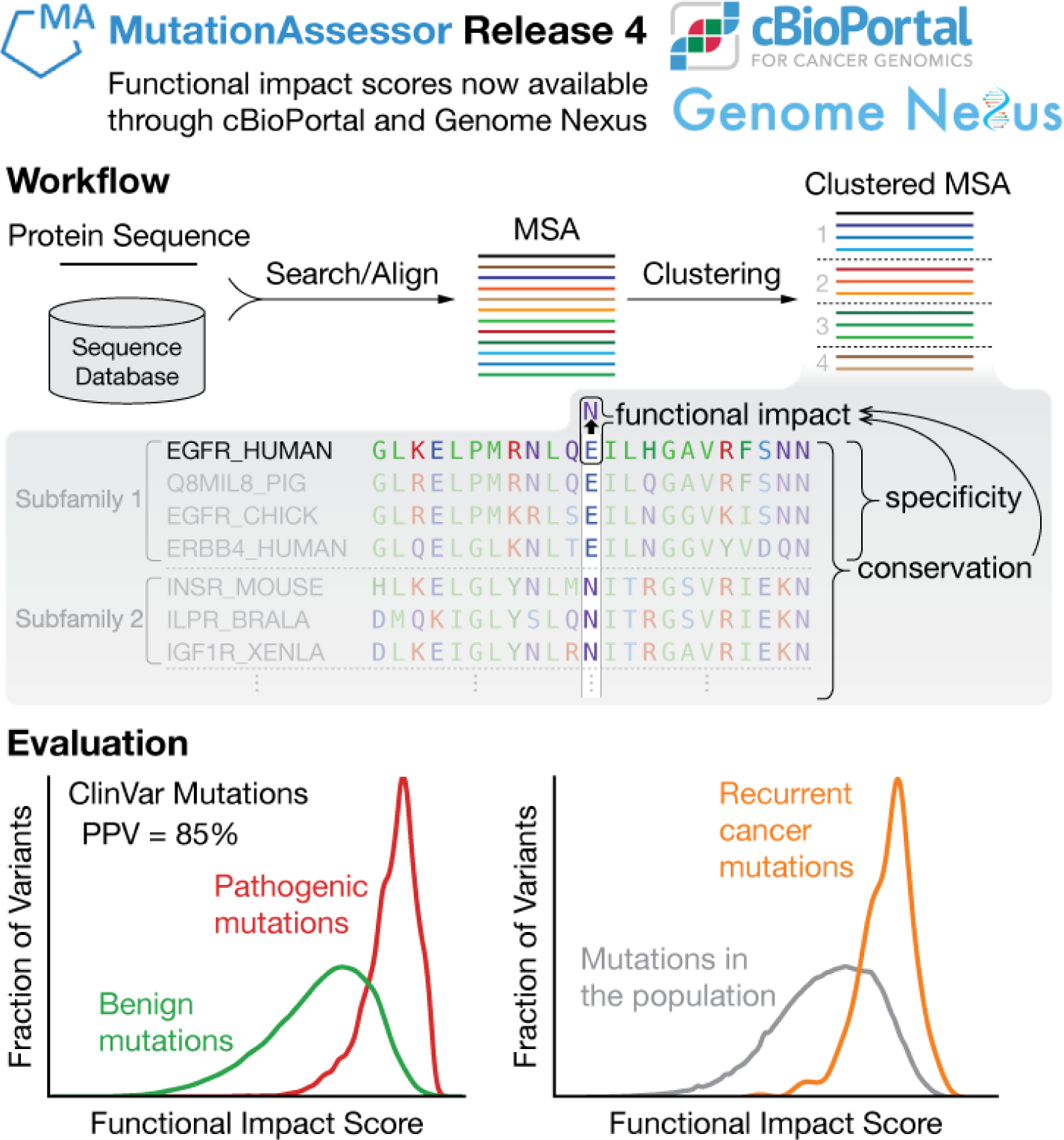
From a mutation in a protein sequence to its functional impact: workflow and evaluation. The functional impact score (FIS) for predicting the impact of a mutation in a protein is derived from the evolutionary conservation pattern of the mutated residue in the protein’s family and subfamily (Methods, Eq. 1–3). Larger FIS indicates more likely adverse functional impact. We use sequences from a homology search to a sequence database to define a protein family. These sequences are aligned to the reference protein and clustered into subfamilies by combinatorial entropy optimization. The relevance of FIS for predicting functional impact can be assessed by its ability to distinguish curated benign and pathogenic mutations from the ClinVar database, and to separate mutations observed in the general population (UK Biobank) from highly recurrent, cancer-tissue somatic mutations (extracted from cBioPortal, number of occurrences in cancer samples > 200).

### Improvement over older version

MA r4 provides updated functional impact scores for all AA-changing single substitution variants in nearly all human proteins, for which multiple sequence alignments are available. This update is based on refined combinatorial entropy analysis of conservation patterns, more comprehensive calculation of multiple sequence alignments covering a larger fraction of human proteins and refined comparison of predicted functional impact with the pathogenic-benign classification in curated knowledge bases such as ClinVar. The updated functional impact scores and multiple sequence alignments are available through [Genome Nexus](https://www.genomenexus.org/) (de Bruijn et al., 2022) and [cBioPortal](https://www.cbioportal.org/) (de Bruijn et al., 2023) (Fig. 2), as well as a downloadable file (DOI: 10.5281/zenodo.15305085).

**Figure 2.**
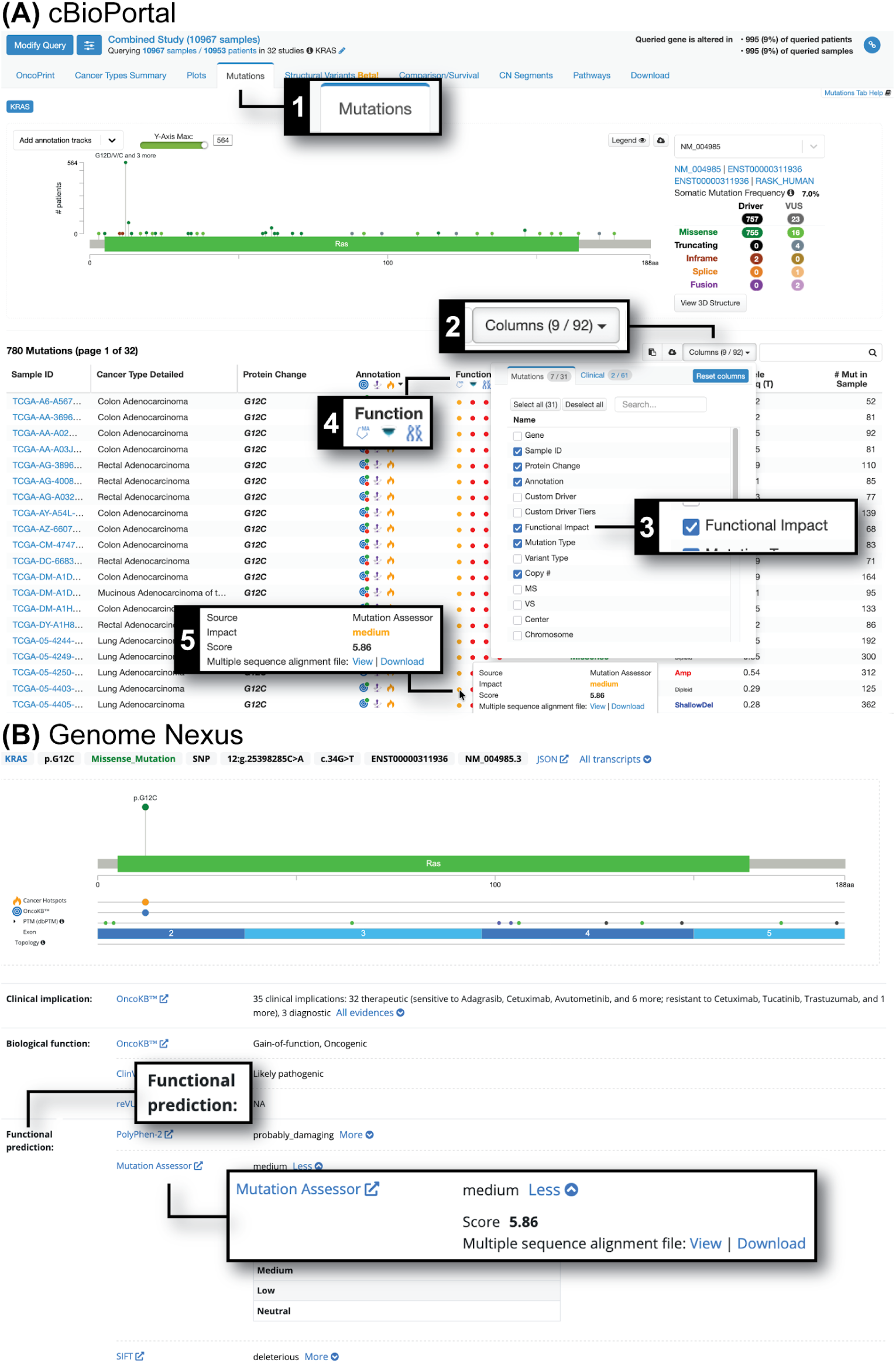
Access to MutationAssessor functional impact predictions in cBioPortal and Genome Nexus. (A) In [cBioPortal](https://www.cbioportal.org/), go to the “Mutations” tab (1), click the “Columns” button (2) and in the pop-up check the “Functional Impact” checkbox (3) to show the “Functional Impact” column in the mutations table. For a particular sequence variant, mouse over the filled circle (5) below the MutationAssessor logo in the “Functional Impact” column (4) to see MutationAssessor predictions. (B) In [Genome Nexus](https://www.genomenexus.org/), MutationAssessor predictions are in the “Functional prediction” section.

### New, deeper alignments of full-length proteins

Functional impact scores (FIS) in MA r4 were computed from newer and deeper multiple sequence alignments (MSAs) built using full-length protein sequence as input to the JackHMMER program (Eddy, 2015) to search and align homologous sequences from the UniRef100 database (Suzek et al., 2015). JackHMMER is based on profile-HMM and is more sensitive for retrieving potentially homologous sequences than BLAST(Camacho et al., 2009) that was used in previous MA releases.

### New, faster CEO implementation

The CEO algorithm that MA uses to cluster the MSAs to identify subfamilies was reimplemented to significantly improve its runtime performance. The new implementation makes heavy use of memoization, sequential memory access, and multithreading. Compared to the single-threaded CEO implementation in our original publication, the new implementation reduced clustering time by ∼16-fold using a single thread or ∼60-fold using 4 threads.

### Selecting FIS cutoffs

As was done in previous MA releases, we assign each AA-changing variant a functional impact based on its FIS in one of four categories: neutral, low, medium and high impact. A higher FIS predicts more likely adverse functional impact. The cutoffs between each category were calibrated against the human-curated clinical labels (benign or pathogenic) in ClinVar (Fig. 3). We chose the cutoff between low and medium impact (FIS = 5.25) such that the false positive rate and false negative rate are balanced, i.e., 82% of the benign variants have a FIS below the cutoff and 82% of the pathogenic variants have a FIS above the cutoff. The cutoff between neutral and low impact was at FIS = 2.6 where 30% of the benign variants and 1% of the pathogenic variants have a FIS below 2.6. The cutoff between medium impact and high impact was at FIS = 7 where 1% of the benign variants and 19% of the pathogenic variants have a FIS above 7 (Fig. 3). A similar analysis for the number of benign and pathogenic variants in the UniProt HUMSAVAR database in each functional impact category is in Supplementary Table S1.

**Figure 3.**
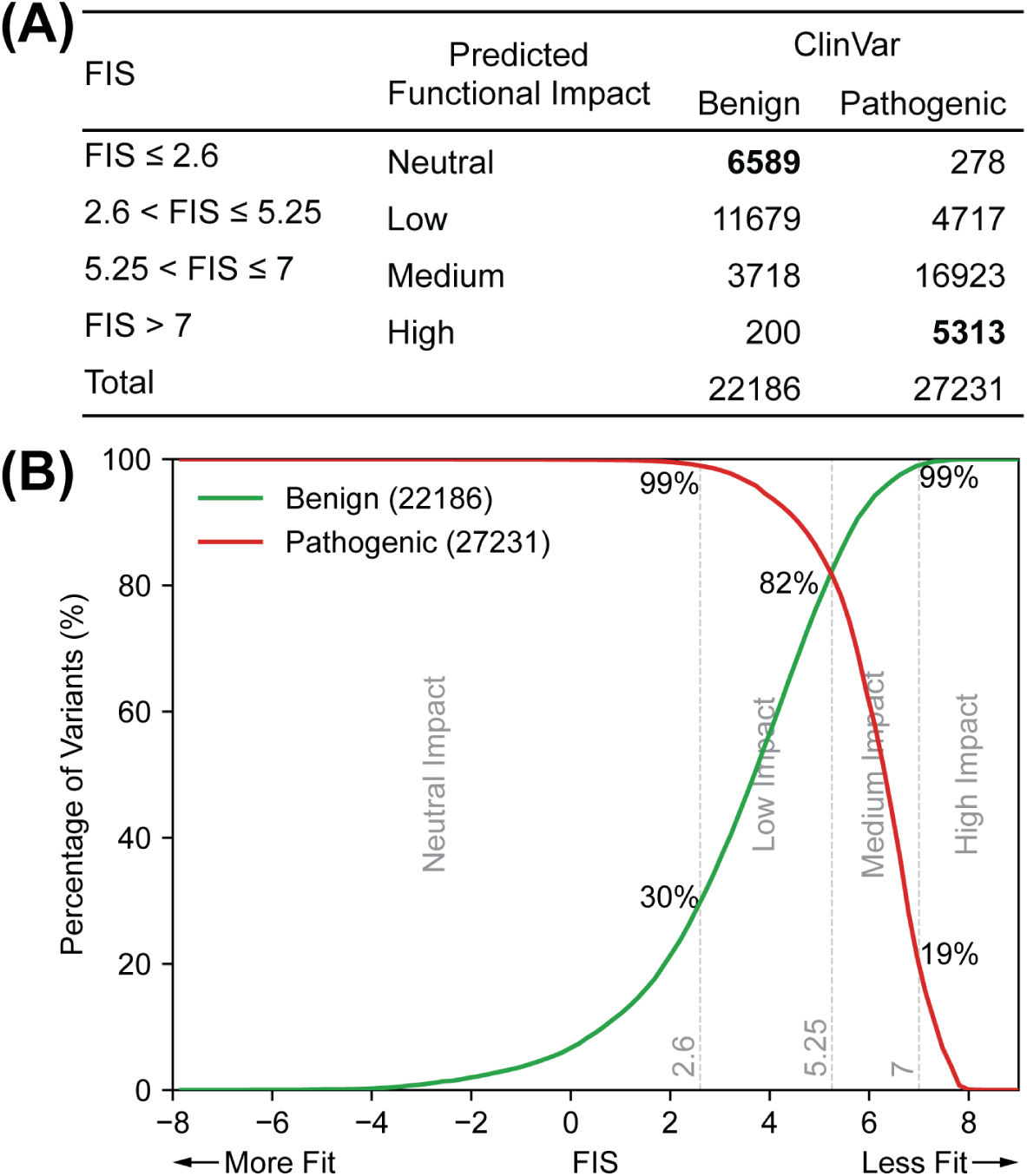
Classification of variant functional impact based on FIS. A variant is assigned to one of four functional impact categories: neutral, low, medium and high impact based on its FIS. **(A)** FIS range for each functional impact category and the number of the benign and pathogenic variants in ClinVar assigned to that category. The vast majority of ClinVar variants that are predicted to be neutral-impact are indeed labelled as benign, and most predicted high-impact variants as pathogenic. The larger values of the bolded numbers indicates good predictive performance. **(B)** FIS distributions of the benign and pathogenic variants in ClinVar. A perfect prediction in a perfect database would have complete separation between the red and green curves.

### Updates with better ClinVar benchmark score

As a strong motivation to update MA functional impact scores in cBioPortal for Cancer Genomics, we found that MA r4 performs better than MA r3 at agreeing with the expert-curated clinical annotations of human disease-related AA-changing single substitution variants from the ClinVar database (Landrum et al., 2014) (average AUC = 0.910 for MA r4 vs 0.877 for MA r3). A caveat is that ClinVar is not necessarily a completely correct gold standard and that it only reports known disease-associated variants. MA r4 also ranks among the top of unsupervised methods evaluated against mutation effects in experimental mutational scans with *in vitro* assays and in ClinVar (Frazer et al., 2021; Notin et al., 2023).

We note that the comparison of MA FIS scores to ClinVar pathogenic-benign annotations only approximately provides an assessment of the quality of the prediction method: ClinVar annotations are not ‘ground truth’ as that database, while highly informative, (1) has acquisition bias and (2) grossly under-reports (for valid practical reasons) benign variants. The best assessment of the quality of the prediction of functional impact would come—and will in part definitely come—from prospective experimental or clinical observation of the impact of variants of currently unknown significance.

### High-FIS variants are rare in the human population

We investigated the population-genetic question to what extent an unsupervised mutation effect analysis and prediction method informed by conservation patterns of individual proteins across many species also reflects selection pressure in the *Homo sapiens* species. Are protein variants deselected in sets of homologs in evolution also underrepresented in the human population?

Without being able to comprehensively analyze this question due to limited data availability, we do observe that single substitution variant frequencies at particular positions in particular proteins in human genomes (UK Biobank 500k Whole Genome Sequencing dataset (Li et al., 2023) and other population variant frequency datasets) and their MA FIS scores are correlated (Fig. 4A, S1, S2).

More frequently observed variants tend to have lower FIS (neutral or low impact) and high-FIS variants are rarely observed in the population. This trend is consistent with the population frequency distributions of ClinVar variants (Fig. 4A bottom), where highly frequent variants in the population are benign (only green bars in the right half) and the pathogenic variants have low frequency in the population (all red bars are in the left half). Our similar analysis using variant frequencies in the NCBI ALFA (Phan et al., 2020) and gnomeAD datasets (Chen et al., 2024) gave a similar relationship (Fig. S1). This result is confirmatory: a simple explanation is that higher mutation impact reflects higher disease propensity leading to deselection in the population; and, that very frequent variants are neutral (aka benign).

**Figure 4.**
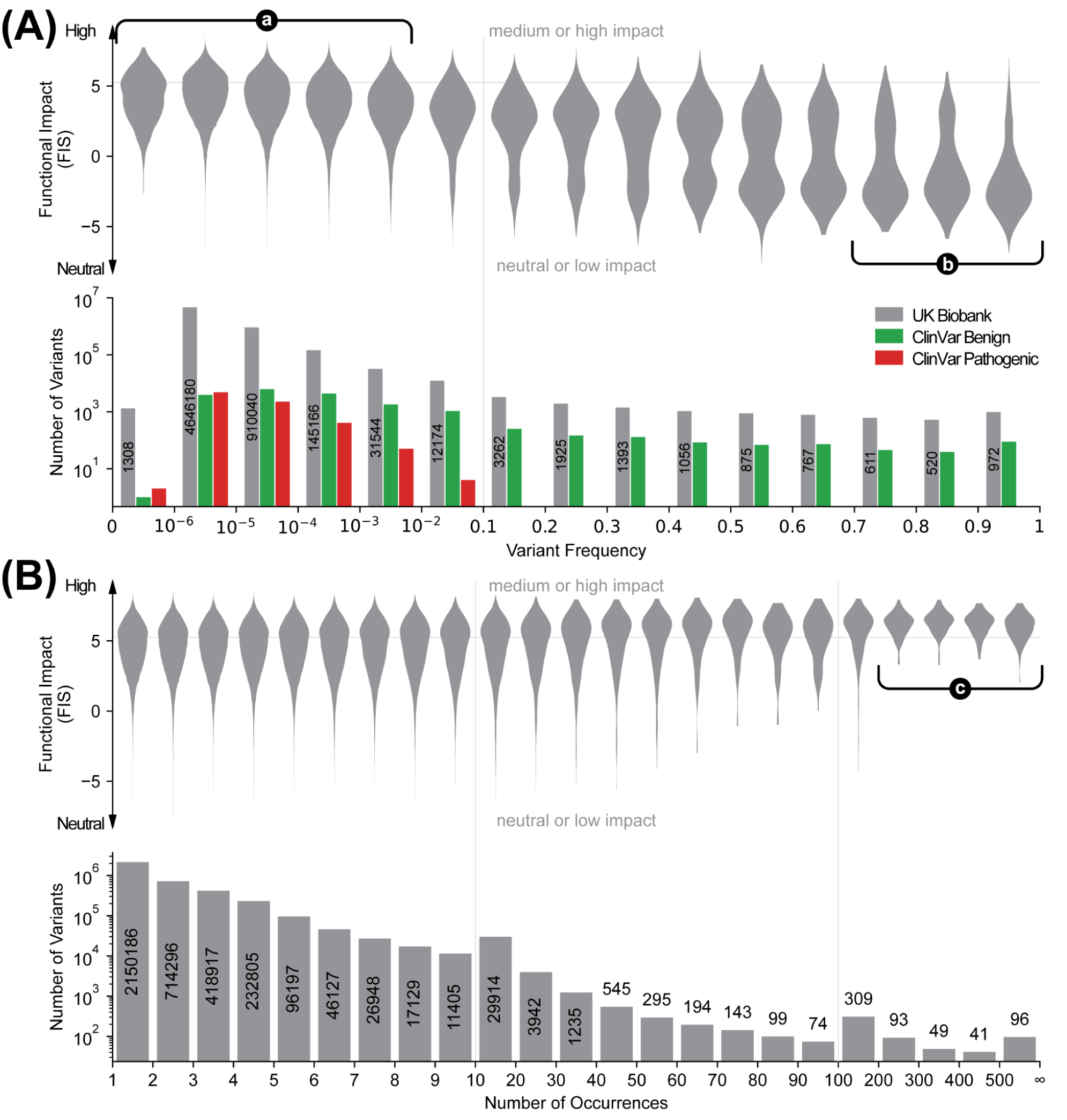
Frequencies and distributions of predicted functional impact of variants in the human population and in cancer tissues. Horizontal lines at FIS = 5.25 indicate the boundary between neutral/low impact and medium/high impact. Vertical lines indicate where the horizontal axis scale changes. **(A)** Functional impact (FIS) and population variant frequencies of AA-changing single substitution variants in the UK Biobank 500k whole genome sequencing dataset. Variants are binned by their frequencies in the population. For each bin, the number of variants therein (lower, gray) and their FIS distribution (upper), the number of variants that are annotated as benign or pathogenic in ClinVar (lower, green or red) are shown. Highlighted regions: (**a**) high-FIS (medium or high impact) variants are rare in the population; (**b**) frequent variants in the population tend to have low FIS (low or neutral impact). (**B**) FIS and number of occurrences of somatic cancer mutations in cBioPortal. Variants are binned by their total number of occurrences across all studies in cBioPortal. For each bin the number of variants therein (lower) and their FIS distribution (upper) are shown. Highlighted region (**c**): highly recurrent variants tend to have high FIS (medium or high impact).

### Recurrent cancer mutations tend to have high FIS

Highly recurrent somatic mutations in cancer tend to have higher FIS. We examined the somatic AA-changing mutations found in cancer patient samples in cBioPortal (Fig. 4B) and COSMIC (Fig. S4) for their number of occurrences and FIS. When mutations are binned by their number of occurrences, high-FIS mutations were particularly enriched in the bins of 200 or more times of occurrences. Of the 279 AA-changing single substitution mutations in those bins, more than 86% are of medium or high impact; in contrast, less than 37% of the mutations that are observed fewer than 200 times are of medium or high impact. Examples (Table 1) of the most frequently observed mutations with predicted high or medium impact include the KRas G12V/C/A/R/S/F mutations that lock it in an constitutively active state due to loss of its GTPase activity (Scheffzek et al., 1997) (G12D being one notable exception as detailed below); the B-Raf V600E/K mutations that make it constitutively active (Wan et al., 2004); the p110α (PIK3CA) E542K, E545K, H1047R mutations that lead to elevated kinase activity (Mandelker et al., 2009; Zhao and Vogt, 2008) (H1047L being another notable exception as detailed below); and the p53 R175H mutation that disrupts folding of its core domain and leading to inactivation (Bullock et al., 2000). However, there were a few cases where highly recurrent oncogenic mutations were predicted to have low or neutral impact. For example, the p110α H1047L (FIS = 2.04) and KRas G12D (FIS = 3.52) mutations were predicted to be neutral and low impact, respectively, despite their well-known association with cancer. The His1047 in WT p110α appeared to be a minority choice, which was reflected by H1047L’s negative conservation score (-1.16) indicating mutating to a more conserved amino acid in the alignment. In the multiple sequence alignment of p110α, only 21% of the sequences had a His at position 1047; the majority of sequences (66%) had Leu (although none of those sequences and p110α belong to the same subfamily). Notably, GEMME (Abakarova et al., 2023) and EVE (Frazer et al., 2021) also predicted those mutations to be more benign than pathogenic among all mutations of the same protein (Supplementary Table S2).

**Table 1.** FIS, GEMME and EVE scores for selected recurrent somatic mutations in cancer tissues. FIS are color coded by their functional impact categories (green: neutral, yellow: low, orange: medium, red: high). To compare FIS, GEMME and EVE scores, which are on different scales, each score’s percentile within the scores of all possible AA-changing single substitution mutations for the same protein are shown in parentheses following the raw scores. Percentile is ranked from least deleterious (0th percentile) to most deleterious (100th percentile).

| Mutant | ClinVar Label | Conservation Score | Specificity Score | FIS | GEMME | EVE |
| --- | --- | --- | --- | --- | --- | --- |
| *KRAS* |  |  |  |  |  |  |
| G12D | Pathogenic | 5.05 | 1.98 | 3.52 (16%) | -1.05 (26%) | 0.62 (59%) |
| G12V | Pathogenic | 6.08 | 5.14 | 5.61 (46%) | -2.98 (68%) | 0.65 (64%) |
| G12C | Pathogenic | 5.89 | 5.83 | 5.86 (50%) | -1.94 (47%) | 0.45 (38%) |
| G13D | Pathogenic | 6.31 | 5.97 | 6.14 (56%) | -1.33 (33%) | 0.44 (37%) |
| G12A | Pathogenic | 3.22 | 5.83 | 4.53 (30%) | -1.09 (27%) | 0.47 (40%) |
| G12R | Pathogenic | 4.79 | 5.83 | 5.31 (41%) | -1.15 (29%) | 0.46 (39%) |
| *BRAF* |  |  |  |  |  |  |
| V600K |  | 8.35 | 6.97 | 7.66 (82%) | -3.26 (70%) | 0.67 (77%) |
| K601E | Pathogenic | 6.45 | 6.97 | 6.71 (49%) | -2.83 (65%) | 0.72 (99%) |
| *PIK3CA* |  |  |  |  |  |  |
| E545K | Pathogenic | 7.09 | 5.26 | 6.17 (72%) | -2.07 (59%) | 0.42 (45%) |
| H1047R | Pathogenic | 5.96 | 4.55 | 5.25 (44%) | -1.94 (56%) | 0.36 (40%) |
| E542K | Pathogenic | 5.93 | 5.26 | 5.59 (52%) | -1.85 (53%) | 0.49 (51%) |
| H1047L | Pathogenic | -1.16 | 5.24 | 2.04 (6%) | -0.96 (24%) | 0.16 (15%) |
| *TP53* |  |  |  |  |  |  |
| R175H | Pathogenic | 6.99 | 4.43 | 5.71 (68%) | -2.84 (65%) | 0.50 (36%) |

### Switch of function variants

Subfamily-specific amino acid conservation may be useful for identifying “switch-of-function” variants, i.e., those that alter the protein’s functional specificity. In the literature, switch-of-function mutations are commonly reported as “gain-of-function”, which is used as an umbrella term that encompasses both mutations that increase the magnitude of the function, but do not alter its nature, and mutations that acquire new functions for the mutant protein. Assuming that “subfamilies” identified by the CEO algorithm have likely distinct functional specificities, an α→β mutation where the wildtype amino acid α and the variant amino acid β are conserved in distinct subfamilies (i.e., they have mutually exclusive conservation patterns) may suggest changed functional specificity for the variant protein relative to the wildtype. Such a mutation will have a high specificity score, typically paired with a low or moderate conservation score, and a high “conjugate specificity score”, which is like specificity score but for the reverse mutation β→α in a subfamily where β is conserved. For example, exchanging a few amino acids between the Ras-related small GTPases RAC1 and CDC42 was sufficient to switch the cell morphology changes induced by them (Heo and Meyer, 2003). All of those switch-of-function mutations have high specificity scores, high conjugate specificity scores and low conservation scores (Table 2). Applying these rules to the highly recurrent mutations in cBioPortal, we identified several putative switch-of-function mutations in cancer (Table 3). Note that not all switch-of-function mutations have a high specificity score and a high conjugate specificity score. As an example, the wildtype IDH1 enzyme converts isocitrate to α-ketoglutarate; the R132C/H/L/S mutations cause changes in the active site and the mutant enzymes instead convert α-ketoglutarate to 2-hydroxyglutarate (Dang et al., 2009). In this case, the R132C/H/L/S mutations all have high specificity scores, but their conjugate specificity scores are very low (negative) and conservation scores are very high (Table 2), indicating that the variant amino acids C/H/L/S are rare or absent at this position in the homologous sequences in the protein family and subfamilies. Indeed, R132 is highly conserved in the multiple sequence alignment.

**Table 2.** Conservation, specificity and conjugate specificity scores of switch-of-function mutations in RAC1, CDC42 and IDH1. All of the switch-of-function mutations in RAC1 and CDC42 (Heo and Meyer, 2003) have low conservation scores, high specificity scores, and high conjugate specificity scores. In contrast, IDH1 switch-of-function mutations have very high conservation scores, high specificity scores and very low conjugate specificity scores. Conjugate specificity scores are computed similarly to the specificity scores but for the variant-to-wildtype reverse mutation in a subfamily where the variant amino acid is conserved.

| Mutation | Conservation Score | Specificity Score | Conjugate Specificity Score |
| --- | --- | --- | --- |
| *RAC1* |  |  |  |
| I33V | -0.89 | 5.12 | 5.64 |
| S41A | 0.92 | 5.12 | 5.53 |
| A42V | 0.66 | 5.12 | 5.96 |
| D47G | 1.54 | 5.12 | 5.54 |
| W56F | 2.08 | 5.12 | 5.16 |
| A95E | -0.69 | 5.11 | 5.63 |
| K116Q | 0.86 | 5.12 | 5.54 |
| A144K | 0.81 | 4.85 | 5.55 |
| R174L | 1.40 | 5.12 | 5.54 |

| *CDC42* |  |  |  |
| --- | --- | --- | --- |
| V33I | 0.69 | 6.19 | 6.09 |
| A41S | -0.92 | 6.20 | 6.33 |
| V42A | -1.07 | 4.81 | 6.18 |
| G47D | -1.37 | 5.50 | 6.33 |
| F56W | -1.92 | 5.51 | 6.33 |
| E95A | 0.79 | 6.19 | 6.08 |
| Q116K | -1.20 | 6.20 | 6.33 |
| K144A | -0.75 | 6.19 | 6.32 |
| L174R | -1.13 | 6.20 | 6.11 |
| *IDH1* |  |  |  |
| R132C | 8.41 | 5.34 | -5.18 |
| R132H | 7.16 | 4.65 | -4.20 |
| R132L | 9.80 | 6.04 | -∞ |
| R132S | 9.80 | 6.04 | -∞ |

**Table 3.** Putative switch-of-function mutations among recurrent cancer mutations. Highly recurrent (≥ 200 occurrences) mutations in cBioPortal that have high specificity scores and high conjugate specificity scores are predicted to be switch-of-function mutations.

| Gene | Mutation | FIS | Conservation Score | Specificity Score | Conjugate Specificity Score |
| --- | --- | --- | --- | --- | --- |
| KRAS | G12A | 4.53 | 3.22 | 5.83 | 5.02 |
| PIK3CA | H1047L | 2.04 | -1.16 | 5.24 | 5.38 |
| ERBB2 (HER2) | R678Q | 3.99 | 2.23 | 5.75 | 4.61 |
| ERBB2 (HER2) | V777L | 3.27 | 0.78 | 5.76 | 4.87 |
| ERBB3 (HER3) | V104L | 3.32 | 1.04 | 5.61 | 4.67 |

## Discussion

### Coverage: full-length versus domains only multiple sequence alignments

In the previous release (MA r3) the multiple sequence alignments were based on protein domains, i.e., subsequences. These were Pfam domains (Paysan-Lafosse et al., 2025) and had overall much shallower MSA depth compared to this release (MA r4) (Fig. 5). We chose to build the new alignments in MA r4 using full-length protein sequences. We hypothesized that for multidomain proteins, alignments of the full-length protein may also capture information about sequence constraints due to inter-domain interactions that would be missed in alignments of individual domains. Thus, FIS scores derived from alignments of full-length proteins more completely reflect the mutational effects on fitness than those derived from separated alignments of domains.

However, this approach excludes conservation information from some shorter sequences in other species that are only alignable to a domain in the human protein, but constraint information from full-length sequences from other species is always included.

**Figure 5.**
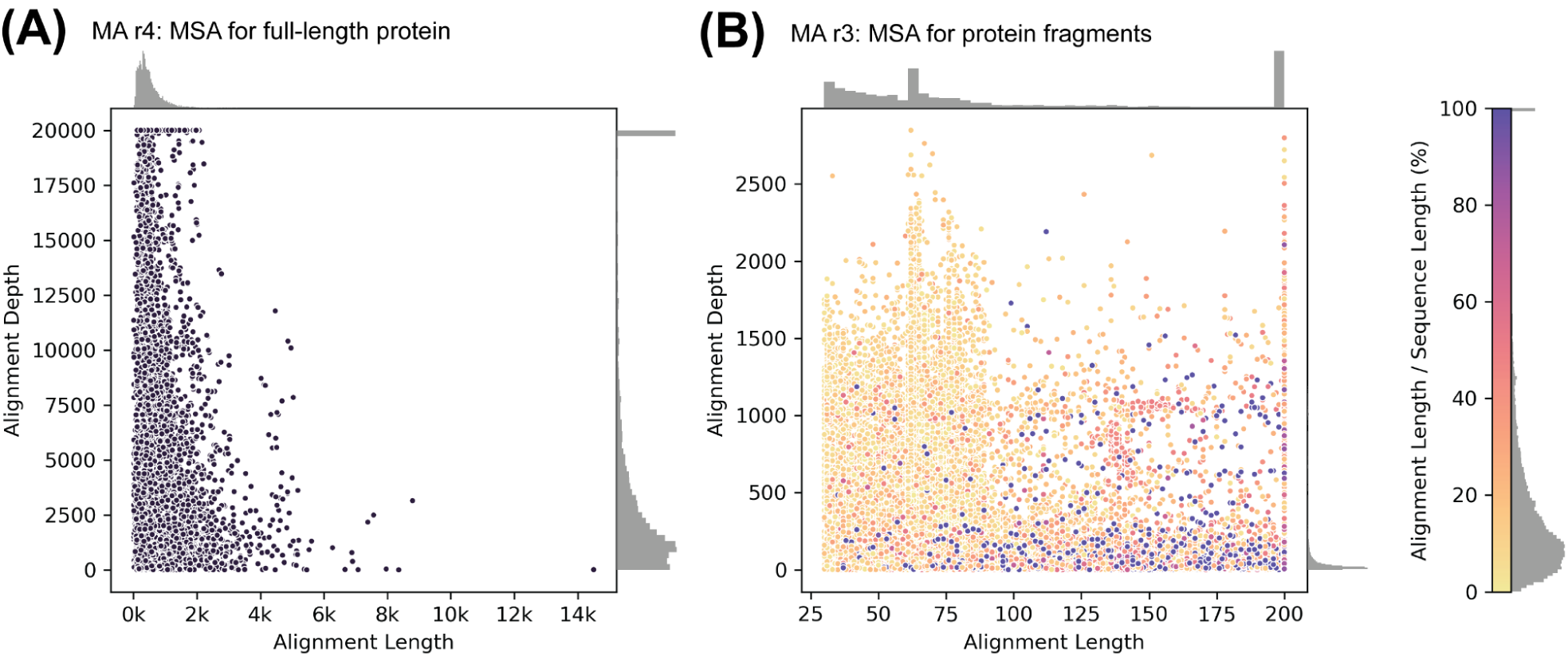
Lengths (number of positions) and depths (number of sequences) of the alignments in MutationAssessor. Each dot represents a multiple sequence alignment of a human protein. Whereas in MA r4 (A) alignments are for full-length protein sequences, in MA r3 (B) they are for protein fragments (often Pfam domains). The fractional lengths of these fragments relative to the full-length proteins range from as low as ∼10% to 100% (color bar). Histograms of length, depth, and fractional length are on the margins.

### Relative information contribution of global conservation and subfamily specificity terms

The predictive power of FIS for the benign and pathogenic clinical labels of the AA-changing single substitution variants in ClinVar is dominated by its conservation score component (Fig. 6). The conservation score clearly provides more discriminative power than the specificity score, as judged by the overlap of the distributions for variants labelled as benign and pathogenic (from human curation). The average AUROC for classifying correctly the benign and pathogenic labels in ClinVar is 0.910 for the combined FIS, 0.906 for the conservation score alone, and 0.812 for the specificity score alone. We explored different weighting schemes of the conservation and specificity score components in FIS and found that approximately equal weighting gave the best classification performance.

**Figure 6.**
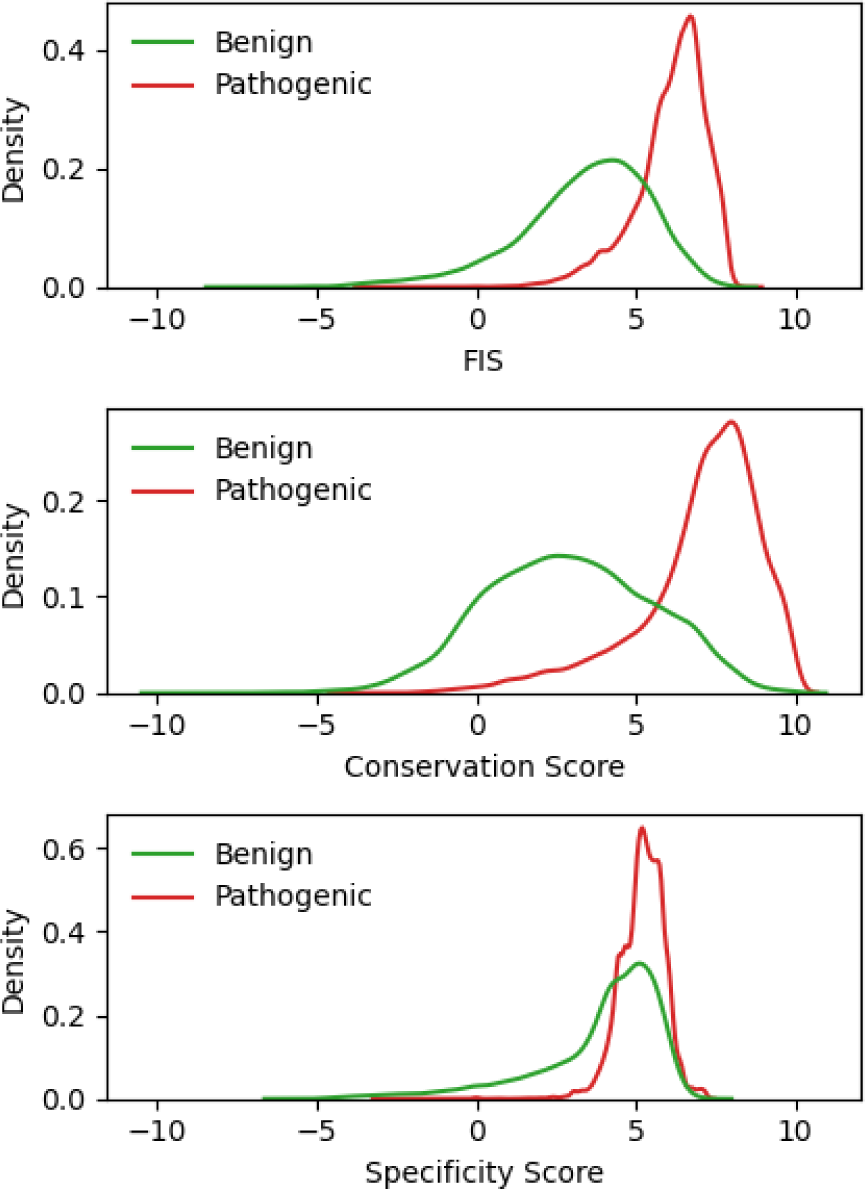
Balance between conservation and specificity scores. The discriminatory power between benign (green) and pathogenic (red) ClinVar single-substitution variants of the combined functional impact score (FIS, top) is mostly due to the conservation score (middle) and less so due to the specificity score (bottom).

### Limitations of the method

MutationAssessor is based on conservation analysis of multiple sequence alignment. As such, its capability is generally limited to amino acid-substitutions at positions where the alignment has good coverage, and is not applicable to other types of sequence variations (e.g., indels or truncations). Due to its use of hierarchical clustering, the CEO algorithm has a time complexity of *O*(*N*^2^ log *N*) where *N* is the depth of the multiple sequence alignment. While our new implementation in MA r4 has vastly improved the runtime efficiency to allow clustering tens of thousands of sequences within a few days, it remains challenging to apply CEO to extremely deep alignments. We are exploring other strategies, including sub-sampling the multiple sequence alignment, ensembling, and alternative clustering algorithms.

### Focus on effects in the human organism versus laboratory based assays

In the landscape of ongoing work about variant effect prediction for different biological questions (Gnad et al., 2013; Notin et al., 2023), the main advantages of this new version of MutationAssessor are its (1) straightforward interpretability in conjunction with (2) improved classification performance for known effects, (3) focus on cancer mutations, and (4) availability of mutation effect predictions in the interactive, user-friendly cBioPortal for Cancer Genomics.

Assessment of prediction performance evaluated on the experimental results of lab-based systematic mutational scans (e.g., all AA variants at all single positions, aka ‘deep mutational scans’) is not the goal of this report. Although not directly relevant, others have made such comparisons (Notin et al., 2023), and MutationAssessor does score reasonably high among unsupervised mutation effect methods in reproducing the impact data from such mutational scans. However, the key caveat is that quantitation of fitness changes in these scans depends on specific experimental assays, which are *in vitro* and narrowly defined and thus of uncertain relevance to *in vivo* consequences for the host organism.

### Recommendations for use by cancer researchers and cell biologists

We suggest that the functional impact scores (FIS) from MutationAssessor are a useful guide for classifying germline and somatic mutations of unknown functional significance, especially in cancer tissues. This information is complementary to assessment of functional roles based on unusually high occurrence frequencies of variants across many cancer samples, relative to random models (often called “significantly mutated”, based on use of a statistical significance threshold (Lawrence et al., 2013). The principal use case is user access to functional assessments of amino acid-substitution variants in specific human genes in one or more tumor types, as a guide to refining functional hypotheses relevant to oncogenesis and therapeutic interventions. When provided by research tools such as the cBioPortal for Cancer Genomics, FIS provides hypotheses that can be useful as a first step in characterizing novel, functional variants and planning focused further experimental investigation on the highest rank candidates using techniques of cell biology including non-human pre-clinical experiments.

## Methods

### Multiple-Sequence Alignments (MSA)

MSAs were generated for all human proteins in the UniProtKB/Swiss-Prot database and selected human proteins in the UniProtKB/TrEMBL database (UniProtKB 2022_02 release) which were referenced in ClinVar (Landrum et al., 2014). For genes that encode multiple protein products, the canonical isoform as specified in UniProtKB was used. Signal peptide (as annotated in UniProtKB) was removed and the resulting sequence was used as the query sequence to perform a series of JackHMMER 3.1b2 (Eddy, 2015) searches using 5 iterations and multiple bit score thresholds against the UniRef100 database. Resulting sequences that align to < 70% of the residues in the query were removed. A range of bit score thresholds were explored until at least 99% of the sequences in the MSA are 30% or more identical to the query sequence.

### Combinatorial Entropy Optimization (CEO)

CEO specificity scoring of residues and sequence clustering (Reva et al., 2007) was performed on MSA columns with at least 70% residue occupancy. At most 20,000 sequences were used for clustering; for larger MSAs, random sampling was used to select subsets of sequences. The *A* parameter was scanned from 0.5 to 0.975 at a step size of 0.025 and the value that maximizes the entropy difference was chosen, resulting in the final choice of granularity of sequence clustering.

### Functional Impact Score (FIS)

After clustering aligned homologous sequences in a protein family into subfamilies by CEO, we compute the FIS (Eq. 1) for a mutation as an average of the conservation score (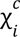, Eq. 2) and and specificity score (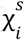, Eq. 3).

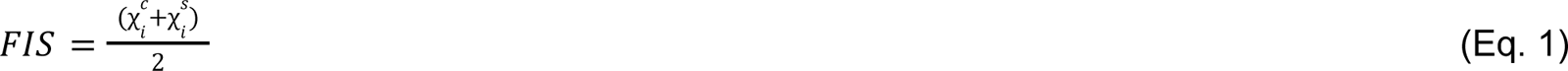

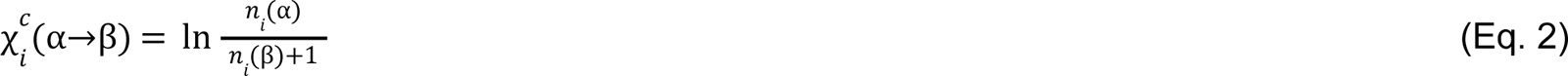

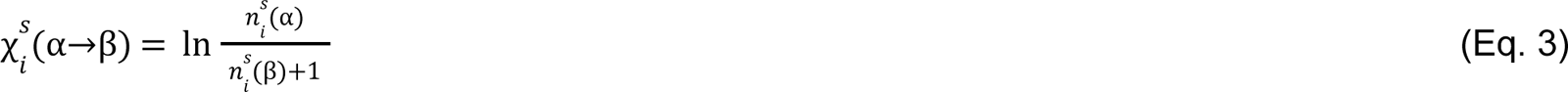

where α and β are the wildtype and mutant amino acids of the mutation, respectively, at the *i*-th sequence position; 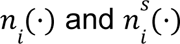 are the number of the specified amino acid at the *i*-th position in the protein family and the subfamily that contains the reference protein, respectively. Note that given a set of amino acids at the *i*-th position of a protein family or subfamily, an α→β mutation increases the number of α by 1 and decreases the number of β by 1. Since the combinatorial entropy is defined as the number of permutations of those amino acids, i.e., 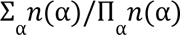 (Reva et al., 2011, 2007), 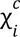 and 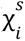 are the changes in combinatorial entropy due to the mutation evaluated in the context of the family and in the context of the subfamily, respectively.

### Variant Frequency

Amino acid level variant frequencies were aggregated from allele frequencies data from UK Biobank 500k Whole Genome Sequencing dataset (Li et al., 2023), NCBI ALFA (Phan et al., 2020), and gnomAD whole genome sequencing dataset v4.0.0 (Chen et al., 2024). Nucleic acid changes were mapped to protein amino acid changes using dbNSFP v4.7a (Liu et al., 2020, 2011). NCBI ALFA and gnomAD allele frequency data were obtained through dbNSFP. Allele frequencies for nucleic acid changes that result in the same amino acid change in the same protein were summed to obtain variant frequencies at the protein level.

### ClinVar

Human AA-changing single substitution variants in the ClinVar database and their expert-curated clinical labels (benign/pathogenic) were obtained following the procedure described in (Frazer et al., 2021). The benchmark dataset contains 30727 benign and 32000 pathogenic variants. Note that not all observed human variants (most of which are plausibly benign) are covered in ClinVar.

### cBioPortal and COSMIC

Cancer somatic mutations were obtained from cBioPortal for Cancer Genomics (including AACR Project GENIE data) and COSMIC Cancer Mutation Census v98.

## Data and Code Availability

The updated functional impact score (FIS) for sequence variants in human proteins and the associated multiple sequence alignments are available by entering the sequence variant at [Genome Nexus](https://www.genomenexus.org/) and [cBioPortal for Cancer Genomics](https://www.cbioportal.org/), and as a file (DOI: 10.5281/zenodo.15305085). Code for the CEO implementation is at https://github.com/sanderlab/ceo. Code for MutationAssessor r4 is at https://github.com/sanderlab/MutationAssessor_r4.

## Acknowledgements

We acknowledge the American Association for Cancer Research for its financial and material support in the development of the AACR Project GENIE registry, as well as members of the consortium for their commitment to data sharing. Interpretations are the responsibility of study authors. Funding for cBioPortal: NCI ITCR 1U24CA274633.

