## Supplementary Materials for "MutationAssessor in cBioPortal"

**Table S1. Predicted functional impact of single AA-substitution mutations in HUMSAVAR.**

| FIS Range | Predicted Functional Impact | HUMSAVAR | |
|---|---|---|---|
| | | Benign | Pathogenic |
| FIS ≤ 2.6 | Neutral | **13977** | 771 |
| 2.6 < FIS ≤ 5.25 | Low | 15398 | 7608 |
| 5.25 < FIS ≤ 7 | Medium | 4646 | 18392 |
| FIS > 7 | High | 315 | **4345** |
| Total | | 34336 | 31116 |

**Table S2. Highly recurrent somatic mutations in cancer.** Somatic mutations that have been observed 300 or more times in total from all studies in cBioPortal are listed. FIS are color coded by their functional impact categories (green: neutral, yellow: low, orange: medium, red: high). GEMME scores and EVE scores for those mutations are included for comparison. Because FIS, GEMME and EVE scores are on different scales, to facilitate comparison, we show each score's percentile within the scores of all possible single AA-substitution mutations for the same protein ranked from least deleterious (0th percentile) to most deleterious (100th percentile).

| Gene | UniProt Accession .Version | Protein Name | Mutant | ClinVar Clinical Label | Conservation Score | Specificity Score | FIS | % | GEMME | % | EVE | % |
|---|---|---|---|---|---|---|---|---|---|---|---|---|
| KRAS | P01116.1 | GTPase KRas | G12D | Pathogenic | 5.05 | 1.98 | 3.52 | 16% | -1.05 | 26% | 0.62 | 59% |
| KRAS | P01116.1 | GTPase KRas | G12V | Pathogenic | 6.08 | 5.14 | 5.61 | 46% | -2.98 | 68% | 0.65 | 64% |
| KRAS | P01116.1 | GTPase KRas | G12C | Pathogenic | 5.89 | 5.83 | 5.86 | 50% | -1.94 | 47% | 0.45 | 38% |
| KRAS | P01116.1 | GTPase KRas | G13D | Pathogenic | 6.31 | 5.97 | 6.14 | 56% | -1.33 | 33% | 0.44 | 37% |
| KRAS | P01116.1 | GTPase KRas | G12A | Pathogenic | 3.22 | 5.83 | 4.53 | 30% | -1.09 | 27% | 0.47 | 40% |
| KRAS | P01116.1 | GTPase KRas | G12R | Pathogenic | 4.79 | 5.83 | 5.31 | 41% | -1.15 | 29% | 0.46 | 39% |
| KRAS | P01116.1 | GTPase KRas | Q61H | Pathogenic | 6.43 | 5.93 | 6.18 | 57% | -1.79 | 44% | 0.64 | 63% |
| KRAS | P01116.1 | GTPase KRas | G12S | Pathogenic | 5.49 | 5.14 | 5.32 | 41% | -1.73 | 42% | 0.30 | 24% |
| KRAS | P01116.1 | GTPase KRas | A146T | Pathogenic | 6.95 | 5.96 | 6.46 | 63% | -1.57 | 38% | 0.46 | 39% |
| KRAS | P01116.1 | GTPase KRas | G13C | Pathogenic | 5.96 | 5.97 | 5.96 | 52% | -1.78 | 43% | 0.31 | 24% |
| KRAS | P01116.1 | GTPase KRas | Q61R | Pathogenic | 5.87 | 5.23 | 5.55 | 45% | -1.96 | 48% | 0.60 | 56% |
| KRAS | P01116.1 | GTPase KRas | Q61L | Pathogenic | 5.74 | 3.22 | 4.48 | 29% | -1.97 | 48% | 0.59 | 53% |
| KRAS | P01116.1 | GTPase KRas | G12F | | 8.35 | 5.83 | 7.09 | 81% | -3.00 | 68% | 0.67 | 68% |
| KRAS | P01116.1 | GTPase KRas | A146V | Pathogenic | 5.72 | 5.96 | 5.84 | 50% | -2.10 | 51% | 0.45 | 38% |
| KRAS | P01116.1 | GTPase KRas | K117N | Pathogenic | 6.77 | 5.97 | 6.37 | 61% | -3.60 | 78% | 0.60 | 55% |
| BRAF | P15056.4 | GTPase KRas | V600E | Pathogenic | 7.26 | 6.97 | 7.11 | 58% | -2.94 | 66% | 0.64 | 73% |
| BRAF | P15056.4 | Serine/threonine-protein kinase B-raf | G469A | Pathogenic | 8.40 | 6.97 | 7.69 | 98% | -3.97 | 79% | 0.71 | 90% |
| BRAF | P15056.4 | Serine/threonine-protein kinase B-raf | V600K | | 8.35 | 6.97 | 7.66 | 82% | -3.26 | 70% | 0.67 | 77% |
| BRAF | P15056.4 | Serine/threonine-protein kinase B-raf | K601E | Pathogenic | 6.45 | 6.97 | 6.71 | 49% | -2.83 | 65% | 0.72 | 99% |
| BRAF | P15056.4 | Serine/threonine-protein kinase B-raf | D594G | | 8.39 | 6.97 | 7.68 | 94% | -5.93 | 93% | 0.72 | 100 % |
| BRAF | P15056.4 | Serine/threonine-protein kinase B-raf | D594N | Pathogenic | 8.39 | 6.97 | 7.68 | 94% | -4.08 | 80% | 0.72 | 99% |
| PIK3CA | P42336.2 | PI3-kinase subunit α | E545K | Pathogenic | 7.09 | 5.26 | 6.17 | 72% | -2.07 | 59% | 0.42 | 45% |
| PIK3CA | P42336.2 | PI3-kinase subunit α | H1047R | Pathogenic | 5.96 | 4.55 | 5.25 | 44% | -1.94 | 56% | 0.36 | 40% |

| | | | | | | | | | | | | |
|---|---|---|---|---|---|---|---|---|---|---|---|---|
| PIK3CA | P42336.2 | PI3-kinase subunit α | E542K | Pathogenic | 5.93 | 5.26 | 5.59 | 52% | -1.85 | 53% | 0.49 | 51% |
| PIK3CA | P42336.2 | PI3-kinase subunit α | R88Q | Pathogenic | 5.72 | 5.26 | 5.49 | 49% | -2.34 | 64% | | |
| PIK3CA | P42336.2 | PI3-kinase subunit α | H1047L | Pathogenic | -1.16 | 5.24 | 2.04 | 6% | -0.96 | 24% | 0.16 | 15% |
| PIK3CA | P42336.2 | PI3-kinase subunit α | N345K | Pathogenic | 6.36 | 5.26 | 5.81 | 57% | -1.63 | 48% | 0.49 | 51% |
| PIK3CA | P42336.2 | PI3-kinase subunit α | C420R | Pathogenic | 6.38 | 5.26 | 5.82 | 57% | -1.34 | 38% | 0.31 | 35% |
| PIK3CA | P42336.2 | PI3-kinase subunit α | E726K | Pathogenic | 3.63 | 5.25 | 4.44 | 27% | -0.93 | 22% | | |
| PIK3CA | P42336.2 | PI3-kinase subunit α | Q546K | | 5.27 | 5.26 | 5.26 | 44% | -2.29 | 63% | 0.55 | 56% |
| PIK3CA | P42336.2 | PI3-kinase subunit α | G118D | Pathogenic | 6.41 | 5.26 | 5.83 | 58% | -1.69 | 49% | | |
| PIK3CA | P42336.2 | PI3-kinase subunit α | E453K | Pathogenic | 7.57 | 5.26 | 6.41 | 79% | -1.97 | 56% | 0.36 | 40% |
| PIK3CA | P42336.2 | PI3-kinase subunit α | Q546R | Pathogenic | 6.52 | 5.26 | 5.89 | 59% | -2.36 | 64% | 0.77 | 72% |
| PIK3CA | P42336.2 | PI3-kinase subunit α | M1043I | Pathogenic | 5.25 | 5.25 | 5.25 | 44% | -1.71 | 50% | 0.08 | 3% |
| PIK3CA | P42336.2 | PI3-kinase subunit α | R108H | | 5.37 | 5.25 | 5.31 | 45% | -1.43 | 41% | | |
| PIK3CA | P42336.2 | PI3-kinase subunit α | E545G | | 7.09 | 5.26 | 6.17 | 72% | -2.40 | 65% | 0.73 | 69% |
| PIK3CA | P42336.2 | PI3-kinase subunit α | E81K | Pathogenic | 6.02 | 5.25 | 5.64 | 53% | -2.87 | 72% | | |
| PIK3CA | P42336.2 | PI3-kinase subunit α | K111E | Pathogenic | 5.24 | 5.26 | 5.25 | 44% | -1.84 | 53% | | |
| PIK3CA | P42336.2 | PI3-kinase subunit α | G1049R | | 5.34 | 5.25 | 5.29 | 45% | -0.93 | 23% | 0.45 | 48% |
| TP53 | P04637.4 | Cellular tumor antigen p53 | R175H | Pathogenic | 6.99 | 4.43 | 5.71 | 68% | -2.84 | 65% | 0.50 | 36% |
| TP53 | P04637.4 | Cellular tumor antigen p53 | R248Q | Pathogenic | 7.21 | 3.28 | 5.25 | 57% | -3.59 | 75% | 0.43 | 29% |
| TP53 | P04637.4 | Cellular tumor antigen p53 | R273C | Pathogenic | 8.60 | 4.41 | 6.50 | 84% | -5.92 | 94% | 0.85 | 73% |
| TP53 | P04637.4 | Cellular tumor antigen p53 | R273H | Pathogenic | 6.65 | 3.31 | 4.98 | 49% | -2.65 | 62% | 0.39 | 26% |
| TP53 | P04637.4 | Cellular tumor antigen p53 | R248W | Pathogenic | 7.50 | 3.28 | 5.39 | 60% | -6.40 | 97% | 0.94 | 88% |
| TP53 | P04637.4 | Cellular tumor antigen p53 | R282W | Pathogenic | 8.59 | 4.43 | 6.51 | 93% | -3.75 | 77% | 0.92 | 83% |
| TP53 | P04637.4 | Cellular tumor antigen p53 | Y220C | Pathogenic | 8.57 | 4.43 | 6.50 | 83% | -3.66 | 75% | 0.88 | 77% |
| TP53 | P04637.4 | Cellular tumor antigen p53 | G245S | Pathogenic | 7.90 | 4.43 | 6.17 | 76% | -3.63 | 75% | 0.63 | 47% |
| TP53 | P04637.4 | Cellular tumor antigen p53 | E285K | Pathogenic | 6.79 | 4.43 | 5.61 | 66% | -4.95 | 87% | 0.87 | 75% |
| TP53 | P04637.4 | Cellular tumor antigen p53 | V157F | Pathogenic | 4.78 | 4.36 | 4.57 | 42% | -2.67 | 63% | 0.86 | 73% |
| TP53 | P04637.4 | Cellular tumor antigen p53 | M237I | Pathogenic | 8.60 | 4.43 | 6.51 | 96% | -2.49 | 60% | 0.81 | 66% |
| TP53 | P04637.4 | Cellular tumor antigen p53 | H179R | Pathogenic | 7.90 | 3.73 | 5.81 | 69% | -6.83 | 99% | 0.96 | 93% |
| TP53 | P04637.4 | Cellular tumor antigen p53 | Y163C | Pathogenic | 8.58 | 4.43 | 6.50 | 86% | -3.69 | 76% | 0.58 | 43% |
| TP53 | P04637.4 | Cellular tumor antigen p53 | S241F | Pathogenic | 7.90 | 3.73 | 5.81 | 69% | -6.83 | 99% | 0.94 | 89% |
| TP53 | P04637.4 | Cellular tumor antigen p53 | R273L | Pathogenic | 8.60 | 4.41 | 6.50 | 84% | -6.09 | 95% | 0.95 | 91% |
| TP53 | P04637.4 | Cellular tumor antigen p53 | R249S | Pathogenic | 8.60 | 4.43 | 6.51 | 97% | -3.78 | 77% | 0.81 | 67% |

| | | | | | | | | | | | | |
|---|---|---|---|---|---|---|---|---|---|---|---|---|
| TP53 | P04637.4 | Cellular tumor antigen p53 | H193R | Pathogenic | 7.90 | 4.43 | 6.17 | 76% | -4.97 | 87% | 0.96 | 93% |
| TP53 | P04637.4 | Cellular tumor antigen p53 | C176F | Pathogenic | 7.90 | 3.73 | 5.81 | 69% | -6.99 | 100 % | 0.96 | 94% |
| TP53 | P04637.4 | Cellular tumor antigen p53 | G245D | Pathogenic | 8.60 | 4.43 | 6.51 | 99% | -6.01 | 94% | 0.94 | 88% |
| TP53 | P04637.4 | Cellular tumor antigen p53 | R158L | Pathogenic | 8.60 | 4.42 | 6.51 | 87% | -6.14 | 95% | 0.88 | 77% |
| TP53 | P04637.4 | Cellular tumor antigen p53 | Y234C | Pathogenic | 8.52 | 4.43 | 6.48 | 80% | -4.17 | 80% | 0.68 | 52% |
| TP53 | P04637.4 | Cellular tumor antigen p53 | I195T | Pathogenic | 8.57 | 4.43 | 6.50 | 83% | -3.97 | 79% | 0.96 | 95% |
| TP53 | P04637.4 | Cellular tumor antigen p53 | H179Y | Pathogenic | 8.60 | 4.42 | 6.51 | 88% | -5.11 | 88% | 0.94 | 88% |
| TP53 | P04637.4 | Cellular tumor antigen p53 | R280T | Pathogenic | 8.60 | 4.43 | 6.51 | 96% | -6.96 | 99% | 0.99 | 100 % |
| TP53 | P04637.4 | Cellular tumor antigen p53 | C238Y | Pathogenic | 8.60 | 4.43 | 6.51 | 98% | -6.78 | 98% | 0.98 | 99% |
| TP53 | P04637.4 | Cellular tumor antigen p53 | P278S | Pathogenic | 8.60 | 4.42 | 6.51 | 89% | -3.43 | 73% | 0.84 | 70% |
| TP53 | P04637.4 | Cellular tumor antigen p53 | E286K | Pathogenic | 7.90 | 4.43 | 6.17 | 76% | -6.19 | 96% | 0.74 | 58% |
| TP53 | P04637.4 | Cellular tumor antigen p53 | R337C | Pathogenic | 4.61 | 4.42 | 4.51 | 41% | -1.11 | 36% | 0.31 | 19% |
| TP53 | P04637.4 | Cellular tumor antigen p53 | R158H | Pathogenic | 6.80 | 3.73 | 5.26 | 57% | -2.70 | 63% | 0.37 | 25% |
| TP53 | P04637.4 | Cellular tumor antigen p53 | Y236C | Pathogenic | 5.96 | 4.33 | 5.14 | 53% | -6.06 | 95% | 0.63 | 47% |
| TP53 | P04637.4 | Cellular tumor antigen p53 | V173L | Pathogenic | 8.60 | 4.43 | 6.51 | 99% | -2.21 | 55% | 0.67 | 51% |
| TP53 | P04637.4 | Cellular tumor antigen p53 | V272M | Pathogenic | 6.90 | 4.43 | 5.67 | 67% | -3.75 | 77% | 0.96 | 94% |
| TP53 | P04637.4 | Cellular tumor antigen p53 | S127F | Pathogenic | 8.60 | 4.43 | 6.51 | 100 % | -6.80 | 98% | 0.87 | 76% |
| TP53 | P04637.4 | Cellular tumor antigen p53 | K132N | Pathogenic | 7.90 | 4.43 | 6.17 | 75% | -4.43 | 83% | 0.84 | 71% |
| TP53 | P04637.4 | Cellular tumor antigen p53 | C176Y | Pathogenic | 8.60 | 4.42 | 6.51 | 88% | -6.99 | 100 % | 0.96 | 95% |
| TP53 | P04637.4 | Cellular tumor antigen p53 | G266R | Pathogenic | 8.60 | 4.43 | 6.51 | 96% | -6.58 | 98% | 0.95 | 91% |
| TP53 | P04637.4 | Cellular tumor antigen p53 | P152L | Pathogenic | 7.90 | 4.43 | 6.17 | 75% | -2.21 | 55% | 0.74 | 58% |
| TP53 | P04637.4 | Cellular tumor antigen p53 | V173M | Pathogenic | 8.60 | 4.43 | 6.51 | 99% | -4.31 | 82% | 0.96 | 93% |
| TP53 | P04637.4 | Cellular tumor antigen p53 | H214R | Pathogenic | 6.32 | 4.43 | 5.37 | 59% | -3.58 | 74% | 0.77 | 63% |
| TP53 | P04637.4 | Cellular tumor antigen p53 | V216M | Pathogenic | 8.58 | 4.43 | 6.51 | 87% | -3.54 | 74% | 0.89 | 78% |
| TP53 | P04637.4 | Cellular tumor antigen p53 | G266E | Pathogenic | 8.60 | 4.43 | 6.51 | 96% | -5.15 | 89% | 0.91 | 81% |
| TP53 | P04637.4 | Cellular tumor antigen p53 | L194R | Pathogenic | 8.60 | 4.43 | 6.51 | 94% | -6.65 | 98% | 0.97 | 97% |
| TP53 | P04637.4 | Cellular tumor antigen p53 | C135Y | Pathogenic | 8.57 | 4.43 | 6.50 | 85% | -6.17 | 95% | 0.97 | 97% |
| TP53 | P04637.4 | Cellular tumor antigen p53 | G245C | Pathogenic | 8.60 | 4.43 | 6.51 | 99% | -5.64 | 92% | 0.95 | 92% |
| TP53 | P04637.4 | Cellular tumor antigen p53 | R280K | Pathogenic | 8.60 | 4.43 | 6.51 | 96% | -2.68 | 63% | 0.96 | 94% |
| TP53 | P04637.4 | Cellular tumor antigen p53 | P151S | Pathogenic | 6.79 | 4.43 | 5.61 | 66% | -2.19 | 55% | 0.77 | 62% |
| TP53 | P04637.4 | Cellular tumor antigen p53 | G245V | Pathogenic | 8.60 | 4.43 | 6.51 | 99% | -6.11 | 95% | 0.96 | 94% |

| | | | | | | | | | | | | |
|---|---|---|---|---|---|---|---|---|---|---|---|---|
| TP53 | P04637.4 | Cellular tumor antigen p53 | K132R | Pathogenic | 8.60 | 4.43 | 6.51 | 97% | -2.46 | 59% | 0.54 | 39% |
| TP53 | P04637.4 | Cellular tumor antigen p53 | C135F | Pathogenic | 8.57 | 4.43 | 6.50 | 85% | -6.17 | 95% | 0.98 | 98% |
| TP53 | P04637.4 | Cellular tumor antigen p53 | C141Y | Pathogenic | 7.18 | 3.73 | 5.45 | 61% | -4.26 | 81% | 0.96 | 94% |
| TP53 | P04637.4 | Cellular tumor antigen p53 | G154V | Pathogenic | 8.57 | 4.43 | 6.50 | 84% | -5.18 | 89% | 0.85 | 73% |
| TP53 | P04637.4 | Cellular tumor antigen p53 | R248L | Pathogenic | 8.60 | 4.38 | 6.49 | 81% | -5.94 | 94% | 0.86 | 74% |
| TP53 | P04637.4 | Cellular tumor antigen p53 | C275Y | Pathogenic | 8.60 | 4.42 | 6.51 | 89% | -6.98 | 99% | 0.98 | 100 % |
| TP53 | P04637.4 | Cellular tumor antigen p53 | Y205C | Pathogenic | 8.59 | 4.43 | 6.51 | 91% | -6.41 | 97% | 0.89 | 79% |
| TP53 | P04637.4 | Cellular tumor antigen p53 | V272L | Pathogenic | 6.90 | 4.43 | 5.67 | 67% | -2.02 | 52% | 0.70 | 54% |
| TP53 | P04637.4 | Cellular tumor antigen p53 | C277F | | 8.60 | 4.43 | 6.51 | 98% | -4.59 | 84% | 0.93 | 87% |
| TP53 | P04637.4 | Cellular tumor antigen p53 | C242F | Pathogenic | 8.60 | 4.43 | 6.51 | 99% | -6.85 | 99% | 0.98 | 100 % |
| TP53 | P04637.4 | Cellular tumor antigen p53 | A159V | Pathogenic | 6.80 | 4.43 | 5.62 | 67% | -2.46 | 59% | 0.40 | 28% |
| TP53 | P04637.4 | Cellular tumor antigen p53 | P278L | Pathogenic | 7.90 | 3.73 | 5.82 | 69% | -5.45 | 90% | 0.77 | 62% |
| TP53 | P04637.4 | Cellular tumor antigen p53 | R110L | Pathogenic | 3.63 | 4.41 | 4.02 | 32% | -1.19 | 38% | 0.62 | 46% |
| TP53 | P04637.4 | Cellular tumor antigen p53 | C238F | Pathogenic | 8.60 | 4.43 | 6.51 | 98% | -6.78 | 98% | 0.97 | 97% |
| TP53 | P04637.4 | Cellular tumor antigen p53 | G266V | Pathogenic | 8.60 | 4.43 | 6.51 | 96% | -6.84 | 99% | 0.97 | 96% |
| TP53 | P04637.4 | Cellular tumor antigen p53 | G244D | Pathogenic | 8.60 | 4.43 | 6.51 | 98% | -3.56 | 74% | 0.70 | 54% |
| TP53 | P04637.4 | Cellular tumor antigen p53 | E271K | Pathogenic | 8.59 | 4.43 | 6.51 | 92% | -3.45 | 73% | 0.79 | 64% |
| TP53 | P04637.4 | Cellular tumor antigen p53 | R213Q | Pathogenic | 7.50 | 3.73 | 5.61 | 66% | -3.32 | 72% | 0.81 | 67% |
| TP53 | P04637.4 | Cellular tumor antigen p53 | E258K | Pathogenic | 8.60 | 4.43 | 6.51 | 97% | -5.53 | 91% | 0.95 | 92% |
| TP53 | P04637.4 | Cellular tumor antigen p53 | C275F | Pathogenic | 8.60 | 4.42 | 6.51 | 89% | -6.98 | 99% | 0.98 | 100 % |
| TP53 | P04637.4 | Cellular tumor antigen p53 | H193Y | Pathogenic | 8.60 | 4.43 | 6.51 | 99% | -4.10 | 80% | 0.89 | 79% |
| TP53 | P04637.4 | Cellular tumor antigen p53 | N239D | Pathogenic | 7.88 | 4.43 | 6.16 | 74% | -3.34 | 72% | 0.93 | 87% |
| TP53 | P04637.4 | Cellular tumor antigen p53 | A161T | Pathogenic | 6.77 | 4.43 | 5.60 | 66% | -2.65 | 62% | 0.89 | 78% |
| IDH1 | O75874.2 | Isocitrate dehydrogenase [NADP] cytoplasmic | R132H | Pathogenic | 7.16 | 4.65 | 5.90 | 36% | -2.14 | 53% | 0.75 | 57% |
| IDH1 | O75874.2 | Isocitrate dehydrogenase [NADP] cytoplasmic | R132C | Pathogenic | 8.41 | 5.34 | 6.88 | 53% | -2.36 | 57% | 0.93 | 97% |
| IDH1 | O75874.2 | Isocitrate dehydrogenase [NADP] cytoplasmic | R132G | | 9.10 | 6.04 | 7.57 | 73% | -4.63 | 89% | 0.92 | 92% |
| IDH1 | O75874.2 | Isocitrate dehydrogenase [NADP] cytoplasmic | R132L | | 9.80 | 6.04 | 7.92 | 93% | -3.97 | 81% | 0.93 | 97% |
| EGFR | P00533.2 | Epidermal growth factor receptor | L858R | Pathogenic | 8.57 | 5.43 | 7.00 | 81% | -5.60 | 92% | 0.85 | 88% |
| EGFR | P00533.2 | Epidermal growth factor receptor | T790M | Pathogenic | 8.80 | 5.43 | 7.12 | 96% | -2.97 | 72% | 0.83 | 81% |
| EGFR | P00533.2 | Epidermal growth factor receptor | A289V | Pathogenic | 8.54 | 5.43 | 6.98 | 79% | -3.86 | 81% | 0.70 | 63% |
| EGFR | P00533.2 | Epidermal growth factor receptor | G719A | Pathogenic | 8.11 | 5.43 | 6.77 | 75% | -3.97 | 82% | 0.86 | 91% |

| | | | | | | | | | | | | |
|---|---|---|---|---|---|---|---|---|---|---|---|---|
| EGFR | P00533.2 | Epidermal growth factor receptor | L861Q | Pathogenic | 8.81 | 5.43 | 7.12 | 99% | -2.05 | 58% | 0.85 | 87% |
| EGFR | P00533.2 | Epidermal growth factor receptor | S768I | Pathogenic | 7.46 | 5.43 | 6.44 | 65% | -3.18 | 74% | 0.77 | 72% |
| JAK2 | O60674.2 | Tyrosine-protein kinase JAK2 | V617F | Pathogenic | 6.64 | 4.73 | 5.69 | 49% | -3.38 | 61% | 0.74 | 65% |
| NRAS | P01111.1 | GTPase NRas | Q61R | Pathogenic | 5.91 | 4.84 | 5.38 | 46% | -1.70 | 49% | 0.60 | 48% |
| NRAS | P01111.1 | GTPase NRas | Q61K | Pathogenic | 6.81 | 4.84 | 5.83 | 54% | -1.91 | 54% | 0.66 | 57% |
| NRAS | P01111.1 | GTPase NRas | G12D | Pathogenic | 5.05 | 4.83 | 4.94 | 38% | -1.11 | 35% | 0.70 | 64% |
| NRAS | P01111.1 | GTPase NRas | Q61L | Pathogenic | 5.77 | 5.54 | 5.65 | 50% | -1.41 | 42% | 0.57 | 45% |
| NRAS | P01111.1 | GTPase NRas | G13D | Pathogenic | 6.31 | 5.54 | 5.92 | 56% | -1.13 | 35% | 0.61 | 49% |
| NRAS | P01111.1 | GTPase NRas | Q61H | | 6.50 | 4.84 | 5.67 | 51% | -1.51 | 45% | 0.64 | 54% |
| NRAS | P01111.1 | GTPase NRas | G13R | | 7.09 | 5.54 | 6.31 | 64% | -2.67 | 68% | 0.53 | 41% |
| SF3B1 | O75533.3 | Splicing factor 3B subunit 1 | K700E | Pathogenic | 7.29 | 6.18 | 6.74 | 51% | -3.09 | 70% | | |
| SF3B1 | O75533.3 | Splicing factor 3B subunit 1 | K666N | | 7.91 | 6.18 | 7.05 | 67% | -4.10 | 84% | | |
| SF3B1 | O75533.3 | Splicing factor 3B subunit 1 | R625C | Pathogenic | 8.00 | 6.18 | 7.09 | 99% | -4.09 | 84% | | |
| U2AF1 | Q01081.3 | Splicing factor U2AF 35 kDa subunit | S34F | Pathogenic | 6.78 | 4.70 | 5.74 | 45% | -2.65 | 64% | 0.90 | 83% |
| U2AF1 | Q01081.3 | Splicing factor U2AF 35 kDa subunit | Q157P | | 5.88 | 5.81 | 5.84 | 47% | -3.01 | 69% | 0.45 | 45% |
| DNMT3A | Q9Y6K1.4 | DNA (cytosine-5)-methyltransferase 3A | R882H | Pathogenic | 8.27 | 5.75 | 7.01 | 75% | -0.57 | 17% | 0.62 | 75% |
| DNMT3A | Q9Y6K1.4 | DNA (cytosine-5)-methyltransferase 3A | R882C | Pathogenic | 8.27 | 5.75 | 7.01 | 75% | -2.07 | 54% | 0.64 | 82% |
| GNAS | P63092.1 | Guanine nucleotide-binding protein G(s) subunit α isoforms short | R201H | Pathogenic | 7.67 | 6.32 | 7.00 | 75% | | | 0.48 | 41% |
| GNAS | P63092.1 | Guanine nucleotide-binding protein G(s) subunit α isoforms short | R201C | | 7.57 | 5.63 | 6.60 | 66% | | | 0.63 | 56% |
| IDH2 | P48735.2 | Isocitrate dehydrogenase [NADP], mitochondrial | R140Q | Pathogenic | 6.89 | 4.81 | 5.85 | 33% | -2.78 | 58% | 0.89 | 65% |
| IDH2 | P48735.2 | Isocitrate dehydrogenase [NADP], mitochondrial | R172K | | 9.09 | 5.51 | 7.30 | 60% | -2.81 | 58% | 0.97 | 97% |
| SRSF2 | Q01130.4 | Serine/arginine-rich splicing factor 2 | P95H | | 4.83 | 4.64 | 4.74 | 44% | -3.72 | 82% | | |
| SRSF2 | Q01130.4 | Serine/arginine-rich splicing factor 2 | P95L | | 4.51 | 4.64 | 4.58 | 39% | -2.61 | 66% | | |
| SRSF2 | Q01130.4 | Serine/arginine-rich splicing factor 2 | P95R | | 2.72 | 4.64 | 3.68 | 17% | -2.67 | 68% | | |
| ERBB2 | P04626.1 | Receptor tyrosine-protein kinase erbB-2 | S310F | Pathogenic | 5.14 | 5.07 | 5.10 | 35% | -2.87 | 68% | | |
| ERBB2 | P04626.1 | Receptor tyrosine-protein kinase erbB-2 | R678Q | | 2.23 | 5.75 | 3.99 | 21% | -0.99 | 29% | | |
| ERBB2 | P04626.1 | Receptor tyrosine-protein kinase erbB-2 | L755S | | 8.59 | 5.76 | 7.17 | 81% | -4.80 | 87% | | |
| ERBB2 | P04626.1 | Receptor tyrosine-protein kinase erbB-2 | V842I | | 5.20 | 5.76 | 5.48 | 41% | -1.86 | 50% | | |
| ERBB2 | P04626.1 | Receptor tyrosine-protein kinase erbB-2 | V777L | | 0.78 | 5.76 | 3.27 | 14% | -1.53 | 42% | | |
| ERBB2 | P04626.1 | Receptor tyrosine-protein kinase erbB-2 | S310Y | Pathogenic | 6.78 | 5.76 | 6.27 | 57% | -2.29 | 59% | | |
| SMAD4 | Q13485.1 | Mothers against decapentaplegic homolog 4 | R361H | Pathogenic | 7.79 | 5.48 | 6.63 | 99% | -2.22 | 53% | 0.71 | 86% |

| | | | | | | | | | | | | |
|---|---|---|---|---|---|---|---|---|---|---|---|---|
| SMAD4 | Q13485.1 | Mothers against decapentaplegic homolog 4 | R361C | Pathogenic | 7.79 | 5.48 | 6.63 | 99% | -3.82 | 75% | 0.73 | 99% |
| FBXW7 | Q969H0.1 | F-box/WD repeat-containing protein 7 | R465C | Pathogenic | 8.17 | 6.83 | 7.50 | 99% | -2.59 | 48% | | |
| FBXW7 | Q969H0.1 | F-box/WD repeat-containing protein 7 | R465H | | 8.17 | 6.83 | 7.50 | 99% | -3.86 | 72% | | |
| FBXW7 | Q969H0.1 | F-box/WD repeat-containing protein 7 | R505C | | 8.16 | 6.83 | 7.50 | 95% | -3.27 | 61% | | |
| FBXW7 | Q969H0.1 | F-box/WD repeat-containing protein 7 | R479Q | | 8.16 | 6.83 | 7.50 | 94% | -3.51 | 66% | | |
| FBXW7 | Q969H0.1 | F-box/WD repeat-containing protein 7 | R505G | | 7.47 | 6.83 | 7.15 | 64% | -4.42 | 80% | | |
| CTNNB1 | P35222.1 | Catenin β-1 | S37F | | 8.09 | 5.92 | 7.01 | 72% | -3.74 | 78% | 0.82 | 83% |
| CTNNB1 | P35222.1 | Catenin β-1 | T41A | | 7.37 | 5.92 | 6.65 | 54% | -2.16 | 53% | 0.78 | 73% |
| CTNNB1 | P35222.1 | Catenin β-1 | S45F | | 6.07 | 5.92 | 6.00 | 35% | -3.60 | 76% | 0.78 | 73% |
| CTNNB1 | P35222.1 | Catenin β-1 | S37C | | 8.09 | 5.92 | 7.01 | 72% | -2.32 | 56% | 0.81 | 80% |
| CTNNB1 | P35222.1 | Catenin β-1 | S45P | | 7.86 | 5.92 | 6.89 | 65% | -1.18 | 33% | 0.73 | 63% |
| CTNNB1 | P35222.1 | Catenin β-1 | S33C | | 7.40 | 5.92 | 6.66 | 54% | -2.39 | 57% | 0.77 | 72% |
| CTNNB1 | P35222.1 | Catenin β-1 | S33F | | 8.09 | 5.92 | 7.01 | 72% | -3.87 | 80% | 0.84 | 90% |
| CTNNB1 | P35222.1 | Catenin β-1 | D32Y | | 8.08 | 5.92 | 7.00 | 71% | -3.84 | 80% | 0.78 | 73% |
| CTNNB1 | P35222.1 | Catenin β-1 | T41I | | 8.06 | 5.92 | 6.99 | 70% | -3.59 | 76% | 0.78 | 73% |
| CTNNB1 | P35222.1 | Catenin β-1 | G34R | | 8.09 | 5.92 | 7.01 | 72% | -4.39 | 88% | 0.82 | 84% |
| BCOR | Q6W2J9.1 | BCL-6 corepressor | N1459S | | 5.36 | 5.07 | 5.21 | 36% | -2.91 | 53% | 0.59 | 76% |
| PTEN | P60484.1 | Phosphatidylinositol 3,4,5-trisphosphate 3-phosphatase and dual-specificity protein phosphatase PTEN | R130Q | | 6.33 | 4.19 | 5.26 | 47% | -4.44 | 90% | 0.73 | 99% |
| PTEN | P60484.1 | Phosphatidylinositol 3,4,5-trisphosphate 3-phosphatase and dual-specificity protein phosphatase PTEN | R130G | | 7.71 | 5.29 | 6.50 | 93% | -7.67 | 100 % | 0.72 | 97% |
| PTEN | P60484.1 | Phosphatidylinositol 3,4,5-trisphosphate 3-phosphatase and dual-specificity protein phosphatase PTEN | R173C | | 7.72 | 5.30 | 6.51 | 99% | -4.97 | 94% | 0.70 | 90% |
| ESR1 | P03372.2 | Estrogen receptor | D538G | | 5.46 | 4.84 | 5.15 | 41% | -4.26 | 81% | 0.55 | 55% |
| ESR1 | P03372.2 | Estrogen receptor | Y537S | | 4.49 | 4.84 | 4.67 | 34% | -2.41 | 49% | 0.50 | 51% |
| CDKN2A | P42771.2 | Cyclin-dependent kinase inhibitor 2A | H83Y | | 7.58 | 4.43 | 6.01 | 98% | -3.99 | 82% | | |
| PPP2R1A | P30153.4 | Serine/threonine-protein phosphatase 2A 65 kDa regulatory subunit A α isoform | P179R | | 7.73 | 4.84 | 6.28 | 66% | -5.60 | 96% | 0.99 | 89% |
| PPP2R1A | P30153.4 | Serine/threonine-protein phosphatase 2A 65 kDa regulatory subunit A α isoform | R183W | Pathogenic | 7.71 | 5.53 | 6.62 | 76% | -4.59 | 90% | 0.99 | 90% |
| PPP2R1A | P30153.4 | Serine/threonine-protein phosphatase 2A 65 kDa regulatory subunit A α isoform | S256F | | 7.72 | 5.53 | 6.63 | 77% | -2.14 | 62% | 0.99 | 87% |
| ERBB3 | P21860.1 | Receptor tyrosine-protein kinase erbB-3 | V104M | | 4.02 | 5.61 | 4.82 | 32% | -0.63 | 15% | 0.35 | 46% |
| FGFR2 | P21802.1 | Fibroblast growth factor receptor 2 | S252W | Pathogenic | 9.38 | 5.25 | 7.32 | 84% | -2.89 | 77% | 0.64 | 67% |

| | | | | | | | | | | | | |
|---|---|---|---|---|---|---|---|---|---|---|---|---|
| FGFR2 | P21802.1 | Fibroblast growth factor receptor 2 | N549K | | 7.84 | 5.25 | 6.54 | 63% | -2.83 | 77% | 0.70 | 72% |
| HRAS | P01112.1 | GTPase HRas | Q61R | Pathogenic | 6.03 | 5.84 | 5.94 | 53% | -2.14 | 52% | 0.61 | 50% |
| SPOP | O43791.1 | Speckle-type POZ protein | F133L | | 3.90 | 5.05 | 4.48 | 19% | -1.04 | 20% | 0.49 | 53% |
| SPOP | O43791.1 | Speckle-type POZ protein | F133V | | 6.08 | 5.05 | 5.56 | 53% | -2.31 | 56% | 0.57 | 69% |
| GNA11 | P29992.2 | Guanine nucleotide-binding protein subunit α-11 | Q209L | Pathogenic | 6.80 | 5.14 | 5.97 | 66% | -3.47 | 81% | 0.84 | 74% |
| H3-3A | P84243.2 | Histone H3.3 | K28M | | 5.20 | 6.12 | 5.66 | 43% | -0.84 | 23% | 0.56 | 33% |
| RAC1 | P63000.1 | Ras-related C3 botulinum toxin substrate 1 | P29S | | 4.55 | 5.12 | 4.84 | 43% | -1.31 | 48% | 0.56 | 32% |
| XPO1 | O14980.1 | Exportin-1 | E571K | | 6.83 | 6.41 | 6.62 | 59% | -3.50 | 71% | | |
| ATM | Q13315.4 | Serine-protein kinase ATM | R337C | | 6.98 | 4.30 | 5.64 | 80% | -5.71 | 95% | 0.84 | 93% |
| KIT | P10721.1 | Mast/stem cell growth factor receptor Kit | V559G | | 7.73 | 5.46 | 6.60 | 74% | -3.16 | 77% | 0.79 | 75% |

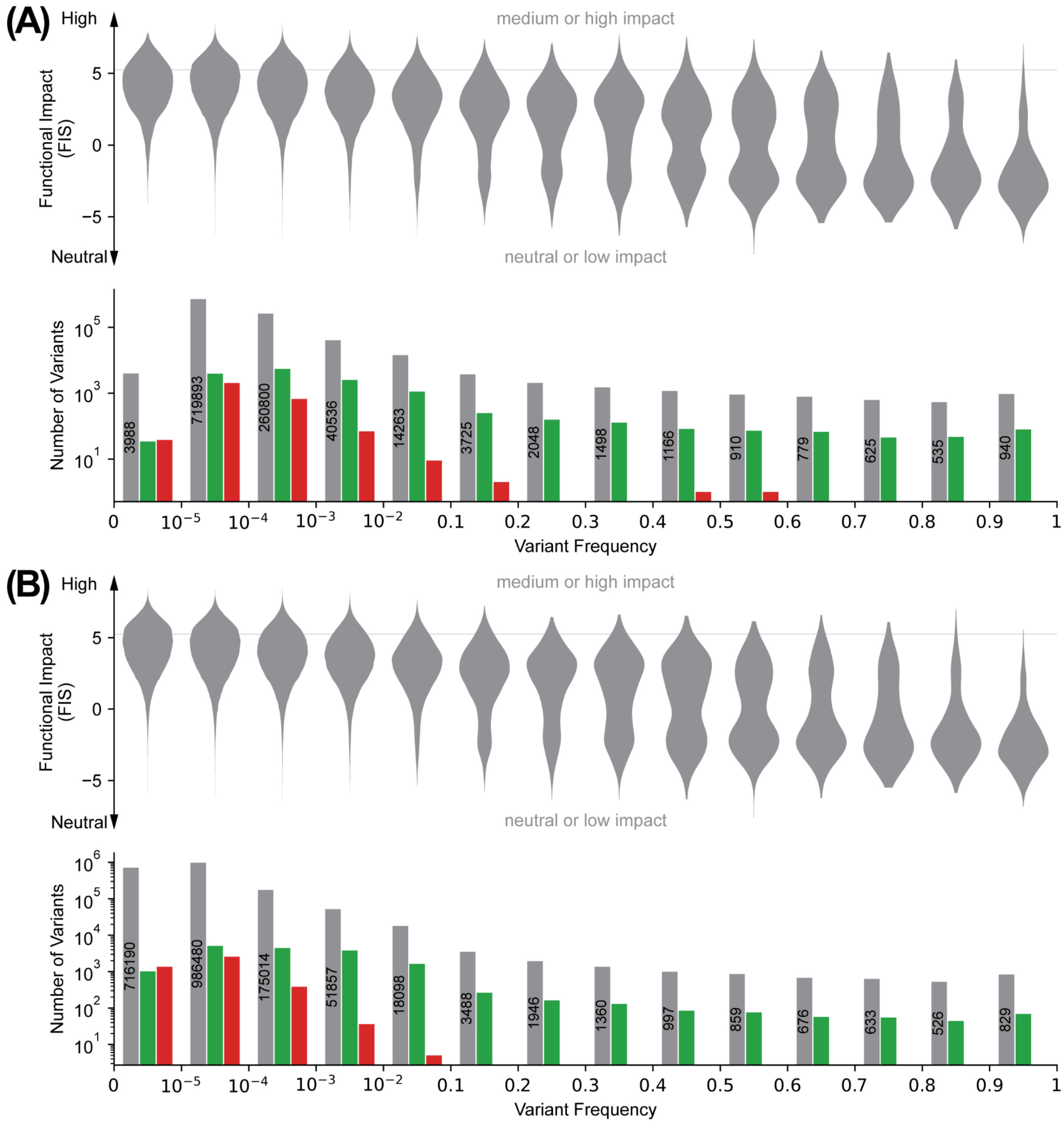


**Figure S1. Frequencies and distributions of predicted functional impact of variants in the population.** Functional impact (FIS) and variant frequencies of AA-changing single substitution variants in the NCBI ALFA dataset (A) and gnomAD whole genome sequencing dataset (B). Variants are binned by their frequencies calculated using the respective dataset. For each bin, the number of variants therein (lower, gray), the number of variants that are annotated as benign or pathogenic in ClinVar (lower, green or red), and their FIS distribution (upper) are shown. Horizontal lines indicate the boundary between neutral/low impact and medium/high impact (FIS = 5.25). Vertical lines indicate where the horizontal axis scale changes.

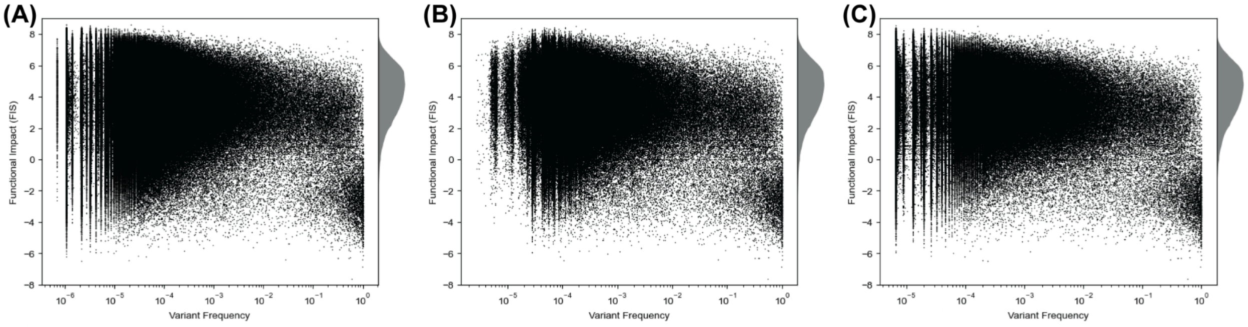


**Figure S2. FIS and variant frequency in the human population.** Each dot represents a unique AA-changing single substitution variant in the UK Biobank 500k whole genome sequencing dataset (A), NCBI ALFA (B), and gnomAD whole genome sequencing dataset (C). A density plot of FIS is shown on the margin of each scatter plot.

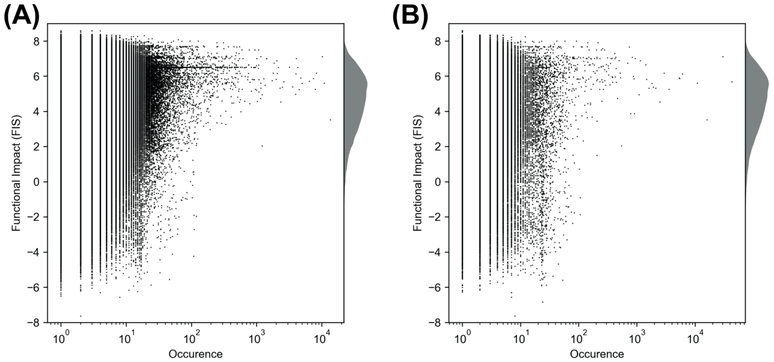


**Figure S3. FIS and the number of occurrences of the somatic variants in cBioPortal (A) and COSMIC (B).** Each dot represents a unique mutation. A density plot of FIS is shown on the margin.

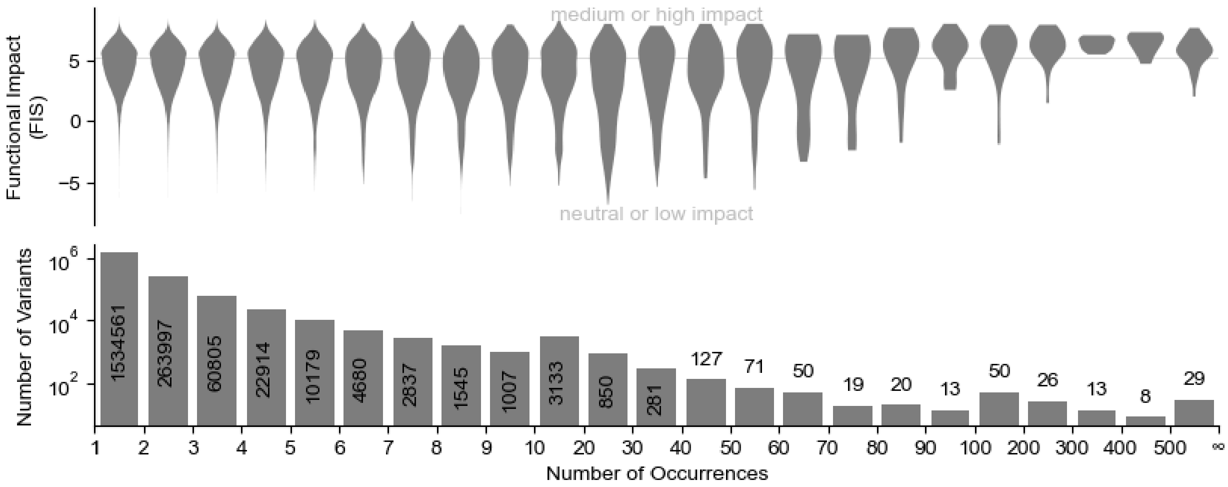


**Figure S4. FIS distribution of cancer mutations from COSMIC Cancer Mutation Census.** Distributions of FIS (**A**) and the number of unique mutations (**B**) stratified by occurrence.
